# Acute degron-mediated RUNX1 loss reprograms enhancer activity to epigenetically drive epithelial destabilization and initiate cancer hallmarks

**DOI:** 10.64898/2026.03.26.711344

**Authors:** Andrew J. Fritz, Haley Greenyer, Louis Dillac, Priyanka Chavarkar, Rahim Ullah, Miles Malik, Deli Hong, Rabail Toor, Emory Pacht, Abigail G. Person, Georgiy Zotkin, Sadie Korzec, Cong Gao, Alquassem Abuorquob, Janine Warren, Jackson Del Porto, Jonah Perelman, Martin Montecino, Jane B. Lian, Andre Van Wijnen, Jessica Heath, Prachi Ghule, Seth Frietze, Kristy Stengel, Scott Hiebert, Kathleen Reed, Tom Misteli, Jonathan A. Gordon, Janet L. Stein, Gary S. Stein

**Affiliations:** Department of Biochemistry, Robert Larner, MD College of Medicine, University of Vermont, Burlington, VT, 05405, USA; National Cancer Institute, National Institutes of Health NIH, Bethesda, MD 20892, USA; Institute of Biomedical Sciences, Faculty of Medicine, Universidad Andres Bello, Santiago, Chile; Millennium Institute Center for Genome Regulation (CRG), Santiago, Chile; Department of Pediatrics, Golisano Children’s Hospital, University of Vermont Medical Center, Burlington, VT, USA; College of Nursing and Health Sciences, University of Vermont Cancer Center, Burlington, Vermont, USA; University of Vermont Cancer Center, Larner College of Medicine, University of Vermont, 89 Beaumont Avenue, Burlington, VT, 05405, USA; Department of Biochemistry, Vanderbilt University School of Medicine, Nashville, TN 37232; The Northern New England Clinical and Translational Research Network, Robert Larner, MD College of Medicine, University of Vermont, Burlington, VT, USA

## Abstract

The RUNX1 transcription factor mediates cell-type specific gene expression. RUNX1 suppression and perturbations are recurrently associated with breast tumor initiation and progression. However, the mechanisms governing the dual roles of RUNX1 in sustaining the mammary epithelial phenotype while epigenetically suppressing initiation of cancer-compromised gene expression are poorly understood. To address this, we used the power of degron-mediated acute, selective, and complete RUNX1 ablation in human mammary epithelial cells. RUNX1 mediates promoter and distal enhancer-driven expression of a gene cohort. Dynamic epigenomic responsiveness upon RUNX1 ablation reveals a rapid and selective decrease in chromatin accessibility and H3K27ac at RUNX1-bound enhancers, but not promoters. While differentially initiated and expressed genes contacted by RUNX1-bound enhancers are enriched in pathways involved in epithelial maintenance and stemness, genes with RUNX1-promoter occupancy support DNA damage responsiveness. Modified cell morphology, metabolic control, increased breast cancer stemness, plasticity, anchorage-independent survival, chemoresistance, and perturbed DNA damage reactivity are observed upon RUNX1 ablation. Together, these findings define RUNX1 as an epigenetic tumor suppressor that maintains epithelial cell state by preserving enhancer activity and preventing gene expression associated with hallmarks of cancer.

## Introduction

Mechanistic and clinical findings are consistent with a role for the RUNX1 transcription factor in supporting breast tissue-specific gene expression as well as implicated in suppressing breast cancer tumorigenicity^1, 2, 3, 4, 5, 6^, epithelial-mesenchymal transition (EMT)^2, 3^, acquisition of phenotype plasticity and stemness^7, 8, 9^. RUNX1 epigenetically mediates transcriptional control of gene expression by forming complexes with the regulatory machinery for histone modifications and chromatin organization^2, 10, 11^. RUNX1 scaffolds these chromatin modulating complexes to mediate physiological responsiveness at multiple regulatory sites of target genes in a context-dependent manner^12, 13, 14^.

There is growing evidence that loss of expression, post-transcriptional regulation, aberrant degradation or inactivating post-translational modifications of RUNX1 may drive epithelial destabilization, cellular plasticity and ultimately, oncogenesis^3, 7, 8, 15^. Functional evidence of RUNX1 involvement in mammary epithelial cell tumor suppression was initially provided by conventional protocols to knockdown transcription factor activity by targeting biosynthesis, stability, or translation of the mRNA^10, 15^. While informative, these knockdown strategies are limited due to reliance on mRNA degradation and persistence of the RUNX1 protein that exhibits delayed turnover (∼8hrs)^16^. These limitations underscore the requirement to define the essential roles of RUNX1 in maintaining the mammary epithelial phenotype. In particular, understanding the immediate, transient, progressive, and sustained consequences of RUNX1 loss is critical for delineating how its absence initiates epigenetic reprogramming and activation of tumorigenic transcriptional programs.

Degron-mediated protein degradation provides a rapid and effective strategy to eliminate transcription factors^10^. The rapid elimination of RUNX1 enables the identification of direct transcriptional and epigenetic consequences. This allows for temporal resolution of both direct and downstream regulatory and phenotypic effects.

We used a degron strategy to address the roles of RUNX1 in both epigenetically mediating regulatory networks that sustain the mammary epithelial phenotype and in enforcing tumor suppressor functions. The effects of RUNX1 loss were studied in normal-like mammary epithelial cells, employing a CRISPR-based approach to tag the endogenous RUNX1 gene with an FKBP12^F36V^ that is ubiquitinated upon the addition of the small PROTAC molecule designated dTAGV1 that recruits the Von Hippel Lindau E3-ligase complex. The addition of dTAGV1 rapidly, selectively and completely degrades RUNX1-FKBP12^F36V^ within 30 minutes. Acute degradation of RUNX1 results in a loss of mammary epithelial cell structure and function, with acquisition of mesenchymal and Breast Cancer Stem Cell (BCSC) phenotypic properties. Anchorage-independent survival, chemoresistance, and perturbed DNA damage responsiveness are observed upon RUNX1 ablation. Breast cancer-compromised gene expression, associated with hallmarks of cancer, is reflected by multi-omic parameters of gene regulation that include transcription initiation (PRO-seq) and RNA expression (RNAseq). RUNX1-dependent epigenetic modifications in the chromatin landscape were reflected by alterations in genome accessibility and organization (ATAC-seq, Micro-C), and genome-wide post-translational histone modifications (ChIP-seq). RUNX1 ablation revealed preferentially decreased chromatin accessibility and H3K27ac at RUNX1-bound enhancers, but not promoters. Our results define a core cohort of RUNX1-responsive genes that support biological control in mammary epithelial cells and provide functional insight into the immediate consequences upon removal of the RUNX1 transcription factor. While differentially initiated and expressed genes contacted by RUNX1-bound enhancers are enriched in pathways involved in epithelial maintenance and stemness, genes with RUNX1-promoter occupancy support DNA damage responsiveness. We conclude that RUNX1 enhancer- and promoter-mediated regulatory network organization and activity reflect epigenetic control of breast cancer tumor suppression.

## Results

### Degron-mediated rapid, selective, and complete ablation of RUNX1 induces a mesenchymal phenotype

We examined RUNX1-mediated epigenetic mechanisms of tumor suppression, and the consequential initiation of breast cancer associated gene expression programs in the absence of RUNX1. We employed a degron system to achieve rapid, selective, and complete degradation of RUNX1 in normal-like mammary epithelial MCF10A cells. This strategy utilizes a RUNX1-FKPB12^F36V^ chimeric knockin in the RUNX1 endogenous locus to inducibly degrade the functional protein by ubiquitination (Figure 1A). Biallelic integration of FKBP at the C-terminus of RUNX1, is confirmed by a mobility shift on western blots, consistent with the incorporation of FKPB12^F36V^ and the absence of wild-type RUNX1 (Figure 1B). This system provides rapid and complete degradation of RUNX1 that is immediately evident as early as 5 minutes and complete by 30 minutes after the addition of the small PROTAC molecule dTAGV1 that recruits the Von Hippel Lindau E3 ubiquitin ligase complex (Figure 1C). dTAGV1-mediated degradation is only observed in degron-containing MCF10A RUNX1-FKBP (MCF10A-R1F) cells but not in parental MCF10A (Figure 1B). To establish that the modified RUNX1-FKBP fusion retains the functional regulatory machinery, coimmunoprecipitation using an HA antibody demonstrates the pulldown of RUNX1, CBFβ, and TLE1 (Figure 1D). These interactions are important as the N-terminal Runt Homology Domain is responsible for DNA binding and CBFβ heterodimer formation, and the C-terminal VWRPY domain is important for interaction with the corepressor TLE1. MCF10A-R1F cells maintain phenotype fidelity of normal mammary epithelial cells that are exhibited by parental unmodified MCF10A^17, 18, 19^. Brightfield microscopic examination demonstrates retention of the normal basal mammary epithelial morphology (Figure 1E, G). Key epithelial markers genes including keratins, CDH1, CDH3, TP63, ACTA2, SMADs, DSP, and COL4A1 are similarly and robustly expressed in MCF10A and MCF10A-R1F cells (Supplementary Figure 1). No significant changes in gene expression are detected upon dTAGV1 treatment in parental MCF10A versus control (DMSO) up to 48hrs indicating selectivity and specificity of the dTAGV1 treatment affecting only modified MCF10A-R1F cells (Supplemental Table 1). RUNX1 ablation by dTAGV1 treatment for 6 days (144hrs) results in less cuboidal cells with more elongated processes indicative of mesenchymal/fibroblastic-like morphology (Figure 1H) that is not observed in parental MCF10A (Figure 1F). This was accompanied by a dramatic reorganization of the actin cytoskeleton as evidenced by F-actin redistribution (Figure 1I,J). These alterations in actin distribution are consistent with changes in cell motility, adhesion, integration with the extracellular matrix, and metastatic potential during breast cancer progression^20^.

**Figure 1.**
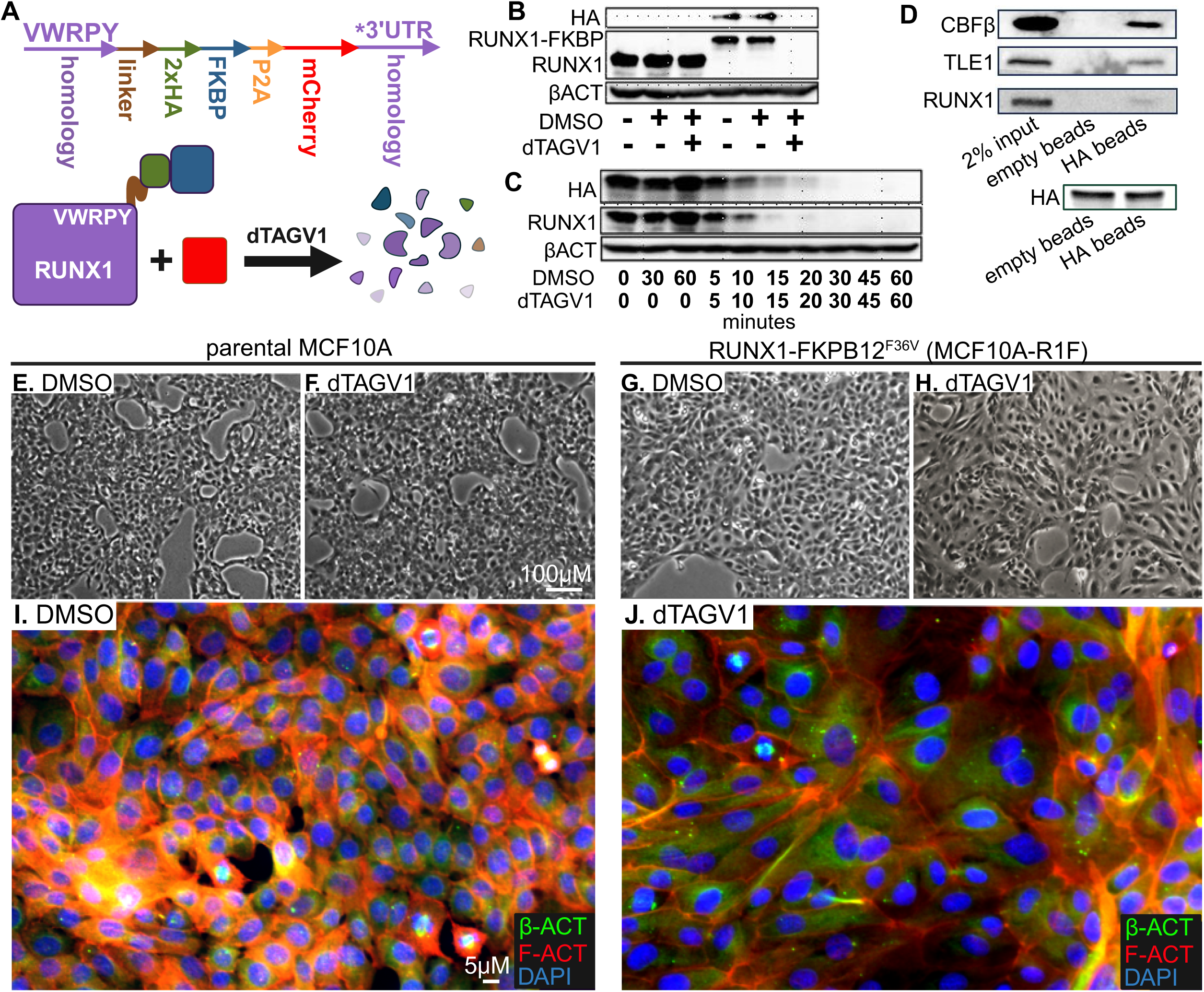
Degron-mediated ablation of RUNX1 results in cytoskeletal reorganization and loss of normal mammary epithelial morphology. A) Schematic illustration of the RUNX1-FKBP degron system. B) Western blot demonstrating a mobility shift consistent with integration of the FKBP epitope and absence of wild type RUNX1 in MCF10A-R1F reflecting biallelic knockin. C) Degradation of RUNX1 is evident by 5 minutes and complete by 30 minutes upon the addition of dTAGV1 PROTAC. D) Immunoprecipitation of FKBP-HA-tagged RUNX1 using an anti-HA antibody demonstrates interaction with CBFβ, TLE1, and modified RUNX1. The same amount of protein was used for the control empty beads and versus the pulldown HA-beads as demonstrated by the same amount of HA detected in both samples. Phase contrast microscopy demonstrating normal epithelial morphology in parental MCF10A E) preceding and F) after dTAG. Normal epithelial morphology in MCF10A-R1F G) prior to RUNX1 ablation is H) altered upon addition of dTAGV1. Immunohistochemistry and fluorescence images depicting Beta Actin and phalloidin of MCF10A-R1F I) without or J) with dTAGV1 treatment.

### Acute RUNX1 depletion rapidly impacts nascent transcription and substantially alters global gene expression

RUNX1 functions as a transcriptional activator or repressor that governs the epithelial phenotype and suppresses tumor initiation^21^. To evaluate the immediate transcriptional response to the loss of RUNX1 in MCF10A-R1F, Differentially Initiated Gene (DIG) transcription was assessed by Precision Run-On sequencing (PRO-seq) at 1hr, 3hrs, and 6hrs. As early as 1hr, there are significant changes (padj <0.05) in transcriptional initiation of 208 genes (log2 fold change ≥0.6 or ≤-0.6) in response to the loss of the RUNX1 protein by dTAGV1 treatment (Figure 2A). At 6hrs, there is a substantial increase in the number of DIGs totaling 2982 genes following RUNX1 ablation. At all timepoints there is a more pronounced downregulation of transcriptional initiation (Figure 2A, Supplementary Figure 2A). PRO-seq enables detection of early transcriptional events that precede changes in steady-state RNA levels (ie. RNA-seq). By RNA-seq there are very few significant (padj>0.05, 8 genes) Differentially Expressed Genes (DEGs) at 1hr. Furthermore, the number of DEGs remains less than the number of detectable DIGs across matched timepoints (Figure 2B). Each PRO-seq timepoint maximally overlaps with a later RNA-seq timepoint. For example, while DIGs at 1hr and 3hrs maximally overlap with DEGs at 12hrs, 6hr DIGs maximally overlap with DEGs at 144hrs (Figure 2C). Notably, ∼50% of DIGs at 6hrs in PRO-seq exhibit corresponding steady-state expression change at later time points in RNA-seq (24hrs and144hrs in RNA-seq; Figure 2C). This lag is consistent with the time required for RNA accumulation to detectable levels.

**Figure 2.**
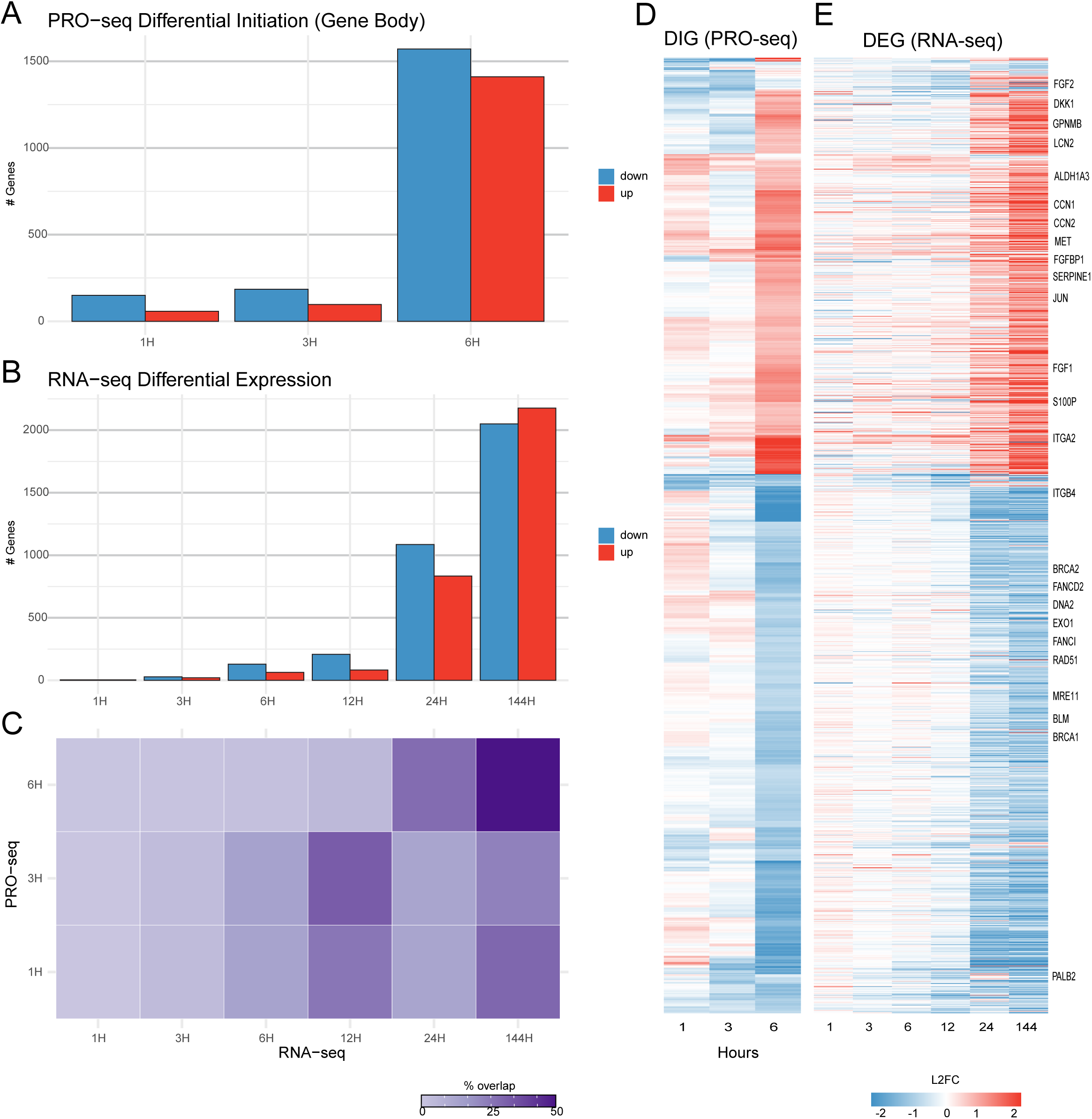
Differential transcriptional initiation and steady-state gene expression upon the loss of RUNX1. A) PRO-seq analysis of DIGs at 1, 3, and 6 hours post RUNX1 ablation (padj <0.05, |log2FC| >0.6) plot indicates the number of significantly differentially initiated genes DMSO vs dTAGV1. B) RNA-seq analysis of DEGs at 1, 3, 6, 12, 24, and 144 hours post RUNX1 ablation (padj <0.05, |log2FC| >0.6) plot indicates the number of significantly differentially expressed genes DMSO vs dTAGV1. C) Heatmap demonstrating the percent correspondence between DIGs and DEGs across measured timepoints following RUNX1 ablation. D,E) Heatmaps depicting the Log2FoldChange of DIGs and corresponding DEGs, respectively. Data represents row normalized z scores. Genes involved in DNA damage response and FGF signaling pathways are highlighted.

Patterns in nascent DIG expression, illustrated by hierarchically clustering, show a distinct transcriptional profile at early timepoints, primarily reflecting the DIGs at 6hrs because this is where most of the genes are differentially initiated (Figure 2D). These dynamic changes reflect an immediate transcriptional response to RUNX1 loss followed by secondary downstream regulatory events. Consistent with gene structural complexity influencing RNA maturation, intron and isoform density are increased with time (Supplementary Figure 2B). Notably, DIG patterns at 6hrs closely resemble DEG profiles observed at 24hrs and 144hrs (Figure 2E). Together, these findings suggest that initial transcriptional events precipitated by the loss of RUNX1 transcriptional activity result in sustained long-term changes in gene expression.

### RUNX1 predominantly regulates cancer-related genes through enhancer-mediated interactions

The DIGs at 6hrs that become DEGs at 24hrs and are sustained up to 144hrs, suggesting that RUNX1 mediates gene expression through direct transcriptional regulation and epigenetic mechanisms. ChIP-seq analysis identifies 22067 sites of RUNX1 genomic occupancy with 4787 of these located within 1kb of transcriptional start sites (TSSs) of DEGs representing promoter proximal binding. These RUNX1 bound promoter proximal sites account for only ∼25% of RUNX1 regulated DEGs, indicating that a substantial portion of RUNX1 activity likely occurs through distal elements such as enhancers. To comprehensively define enhancers in MCF10A, we applied chromHMM analysis, integrating multiple histone post-translational modifications. Enhancer states were primarily defined by H3K4me1 and H3K27ac (Supplemental Figure 3) and further refined by intersecting with ENCODE defined cis-regulatory elements (CREs) from normal breast tissue (Supplemental Table 1). Of 75264 putative identified enhancers, 8913 are bound by RUNX1 (Supplemental Table 1). To map RUNX1-bound enhancers to their target genes, we applied Activity-By-Contact (ABC) modeling in MCF10A cells. ABC integrates each enhancer’s chromatin accessibility, gene expression, and 3D contact frequency from micro-C analysis with candidate promoters to compute an activity-by-contact score, which predicts enhancer–promoter interactions. This subset of RUNX1-bound enhancers predicted by ABC to have high contact frequency with promoters of genes that are differentially initiated and expressed (n = 391) provides a basis for subsequent analyzing the network of RUNX1-enhancer mediated gene regulation.

To directly assess changes in chromatin accessibility and H3K27ac histone post-translation modification at RUNX1-bound enhancers following RUNX1 degradation, we performed ATAC-seq and ChIP-seq, respectively (Figure 3A). RUNX1-bound enhancers exhibited a striking decrease in both ATAC and H3K27ac signals over time, particularly at 144hrs. In contrast, a matching set of non-RUNX1-bound enhancers do not show a comparable reduction in ATAC signal. Moreover, H3K27ac signal at a matched random set of non-RUNX1-bound enhancers is consistently lower than RUNX1-bound enhancers suggesting that RUNX1-bound enhancers are actively marked in the normal mammary epithelial state (i.e. control) and are dynamically modified during the transition induced by acute RUNX1 loss.

**Figure 3.**
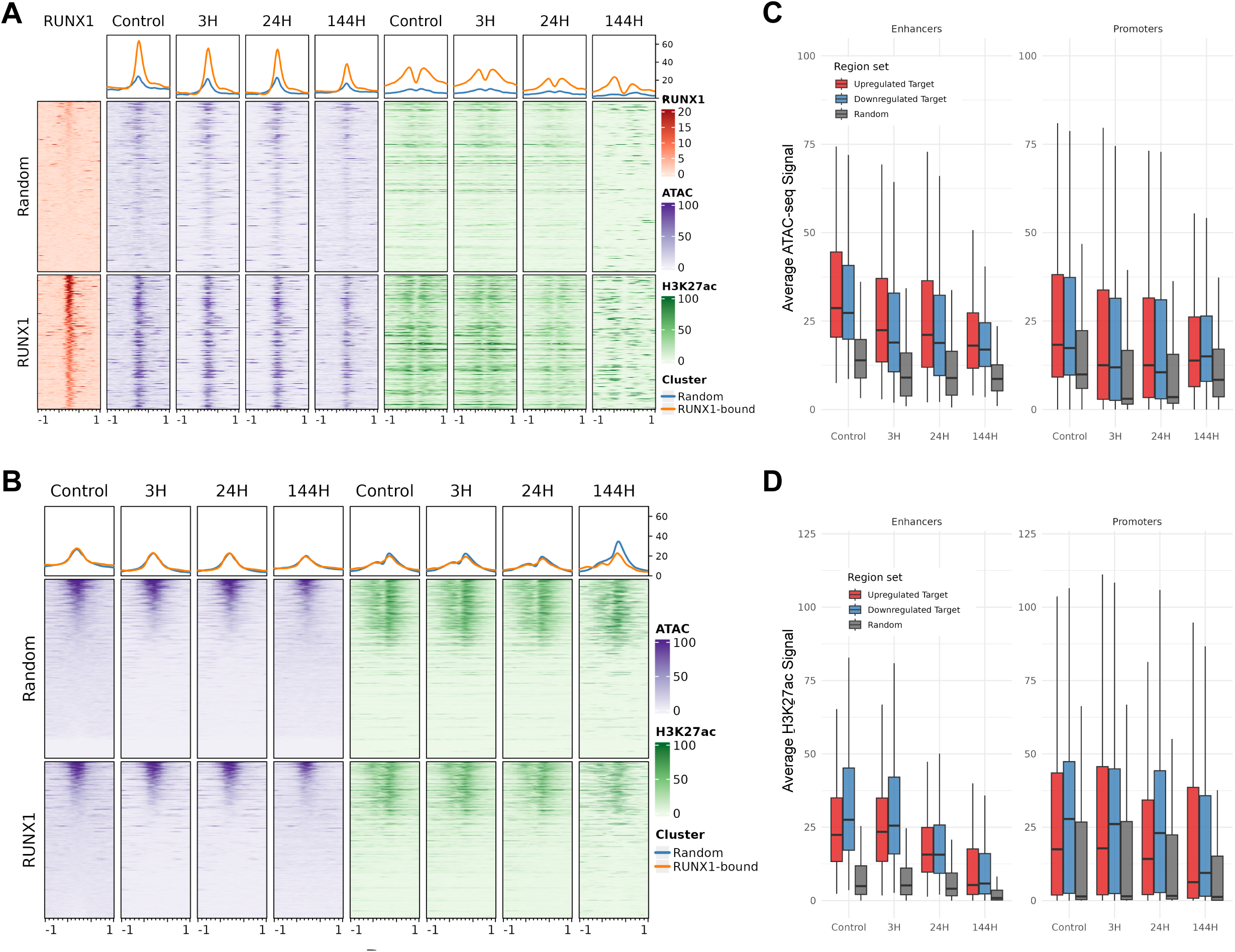
Dynamic remodeling of ATAC and H3K27ac signal at distal enhancers and promoters upon loss of RUNX1. A) ATAC-seq and H3K27ac ChIP-seq signal heatmaps binned over RUNX1-bound DIG/DEG promoters and randomly selected non-RUNX1-bound promoters across time (see Methods). Aggregate line plots (top) summarize temporal trends exhibited by the two promoter sets. B) ATAC-seq and H3K27ac ChIP-seq signal heatmaps binned over both RUNX1-bound distal enhancers contacting DEGs and randomly selected non-RUNX1-bound enhancers across time. Aggregate line plots (top) summarize temporal trends exhibited by the two enhancer sets. C) Boxplots illustrate average ATAC signal at enhancers and promoters across time, grouped by the expression status of associated genes (upregulated, downregulated, or random). D) Boxplots illustrate average H3K27ac signal at enhancers and promoters across time, grouped by the expression status of associated genes (upregulated, downregulated, or random).

In contrast to enhancers, RUNX1-bound promoters associated with DEGs show minimal changes in ATAC and H3K27ac signal over time and signal intensity is similar to that observed a matched random set of non-RUNX1-bound promoters (Figure 3B). Notably, RUNX1-bound enhancers linked to DEGs exhibit a greater reduction in mean ATAC signal regardless of whether the associated gene was up or downregulated. In comparison, the mean ATAC signal at random enhancers is consistently lower at all timepoint and remains unchanged over time (Figure 3C). RUNX1-bound gene promoters do not show the same change in average ATAC mean signal intensity suggesting that changes in chromatin accessibility at enhancers is more dynamic upon RUNX1 ablation. H3K27ac largely demonstrates the same decrease in average signal intensity as a function of time at enhancers (Figure 3D). This reduction in signal intensity is not observed at gene promoters. Overall, these results indicate that enhancers are more dynamic in response to the loss of RUNX1 than promoters and mediate cancer-related gene expression changes.

### RUNX1 ablation reprograms mammary epithelial gene expression toward cancer-associated transcriptional signatures

The large number of differentially initiated and expressed genes (DIGs and DEGs, respectively) upon RUNX1 loss implicates broad regulatory changes. ClueGO functional network analysis of the union set of DIGs and DEGs revealed activation of signaling pathways including FGF, WNT, TGFβ, and ERK, partial epithelial-mesenchymal transition (hybrid EMT), extracellular matrix (ECM) remodeling, and invasion, alongside repression of DNA damage repair and cell cycle checkpoint pathways (Figure S4A). Representative enhancer-regulated targets include upregulation of *FGF2*, which is upregulated following RUNX1 loss (Figure 4A), and and *DMBT1*, which is downregulated (Figure 4B). In contrast, ClueGO analysis of promoter-driven DEGs indicated consistent downregulation of genes associated with DNA replication, repair, and checkpoint control (Figure 4C). Reflecting the observation that enhancers show greater chromatin remodeling than promoters after RUNX1 degradation, ClueGO analysis of enhancer-linked targets demonstrated significant upregulation of FGF-JUN-ERK signaling (Figure 4D). These pathway-level changes were further supported by individual gene expression shifts, including activation of FGF signaling and repression of DNA damage repair regulators (Figure 4E, F). Together, these findings suggest that RUNX1 loss drives transcriptional reprogramming that promotes epithelial plasticity, impairs genome stability, and enhances signaling pathways associated with transformation and stemness.

**Figure 4.**
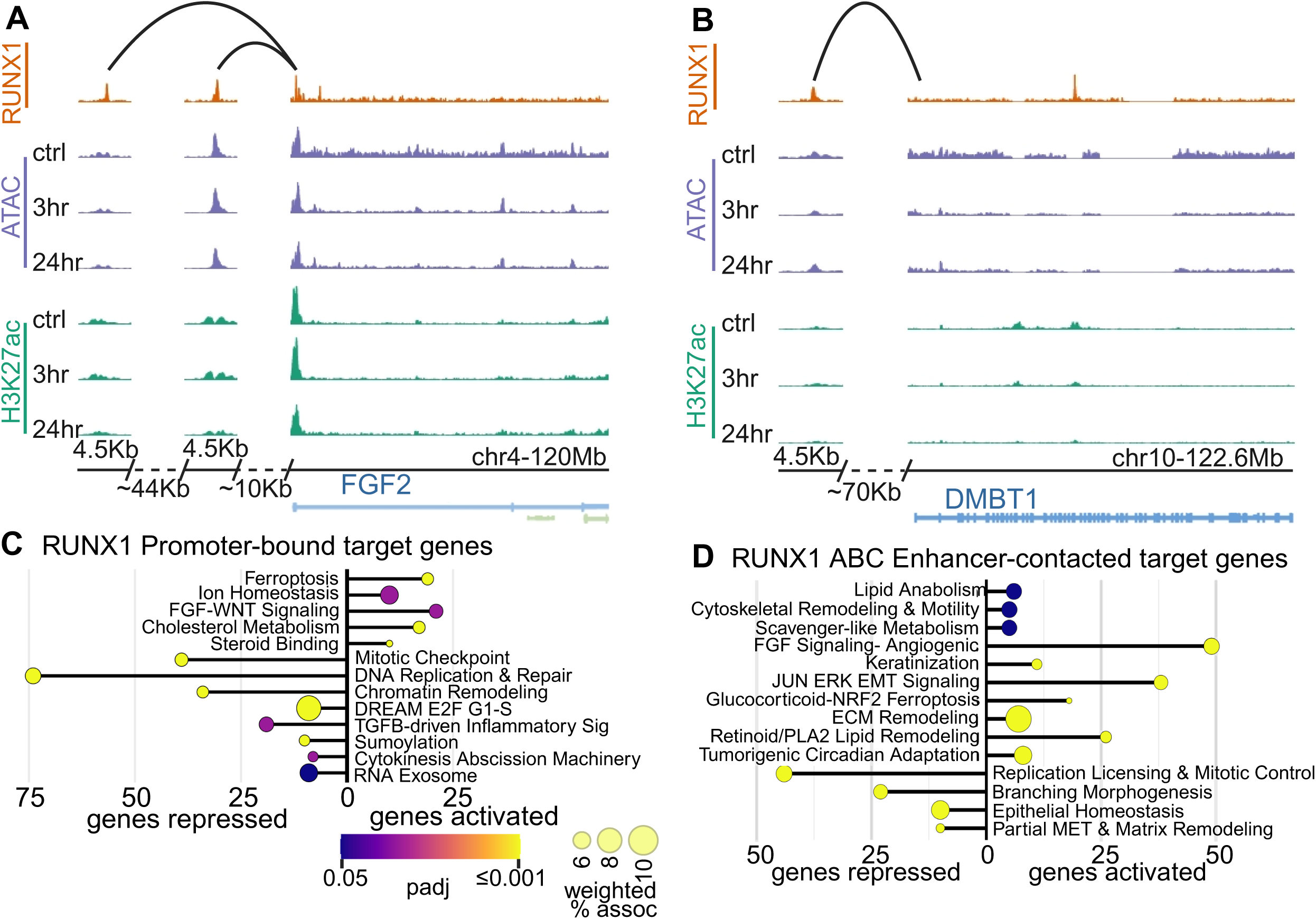
RUNX1-bound distal enhancers or promoters regulate gene expression associated with pathways linked to hallmarks of cancer. Chromosomal tracks depicting RUNX1 binding, chromatin contacts, ATAC and H3K27ac signals at A) chr4-120Mb region encompassing the FGF2 or B) the chr10-122Mb DMBT1 loci. Both genes are regulated through enhancer contact and differentially regulated. C) Functionally grouped network ClueGo analysis for gene promoters that are bound by RUNX1 and DIG/ DEG upon RUNX1 ablation. D) Functionally grouped network ClueGo analysis for RUNX1-bound distal enhancer-associated DIG/ DEGs upon RUNX1 ablation. Color scale indicates padj values and bubble size indicates weighted percent association defined pathways. Heatmap of row normalized z-score values of select genes associated with E) FGF signaling and F) DNA damage response.

### RUNX1 modulates endogenous FGF and TGFβ signaling in the absence of additional exogenous cytokines

The immediate responsiveness of the RUNX1 degron system revealed early enrichment of FGF pathway genes, as shown by ClueGo functional network analysis (Figure 4). This included upregulation of multiple secreted ligands such as FGF2, FGFBP1, BMP2, CCN1, and CCN2, all of which showed increased transcriptional initiation and steady-state mRNA expression. Notably, RUNX2, a paralog of RUNX1 and a known oncogenic driver in advanced breast cancer^5^, was minimally expressed in MCF10A cells under basal conditions. Upon RUNX1 degradation, RUNX2 was transcriptionally induced as early as 3 hours, with RNA accumulation preceding a detectable increase in protein by 12 hours (Supplementary Figure 5). This delayed protein upregulation highlights a reciprocal relationship between the RUNX1 tumor suppressor and the RUNX2 oncoprotein. Given their shared DNA binding domains of RUNX1 and RUNX2, this finding supports potential competition at *cis*-regulatory elements controlling genes involved in epithelial transformation, but as a secondary event after 12 hours. The delayed upregulation of RUNX2 at the protein level supports RUNX1-mediated regulation of FGF signaling that is not responsive to RUNX2 during the initial 12 hours following RUNX1 loss.

In parallel, TGFβ pathway components were dynamically reprogrammed. Transcription of TGFβR2 was transiently downregulated, while TGFβR3 expression increased, a shift consistent with prior results implicating RUNX1 as a direct regulator of TGFβR3^7^. Since TGFβR3 modulates non-canonical TGFβ signaling through SMAD1/5/9, whereas TGFβR2 acts via SMADs 2/3^22^, this receptor switch suggests a transition towards alternative TGFβ and BMP signaling following RUNX1 loss. TGFβR3-regulated targets were significantly enriched at 12 hours post-degradation. These transcriptional shifts were consistent with activation of downstream effectors, as both MAPK and PI3K signaling cascades were increasingly activated at 24 and 144 hours. Importantly, these changes occurred without the addition of exogenous FGF, BMP, or TGFβ ligands, demonstrating that RUNX1 functions upstream of these signaling programs, acting as a key repressor of intrinsic epithelial cytokine signaling that becomes unleashed upon its loss.

### RUNX1 degradation promotes Breast Cancer Stem Cell (BCSC) emergence, anchorage-independent survival and chemoresistance

Aldehyde dehydrogenase activity of BCSCs is largely attributable to the ALDH1A3 isoform, and its expression predicts metastasis^23^. We identified ALDH1A3 as a critical target of RUNX1-bound enhancers from our enhancer analysis. ALDH1A3 is contacted by two small (349, 213bp) CREs that are located ∼8kb and ∼167kb from the gene TSS (Figure 5A). These enhancers, although small, were associated with strong ATAC and H3K27ac signals that are broader than the identified enhancer. This interaction between these RUNX1 bound enhancers and the gene promoter suggests that RUNX1 regulates this gene through an enhancer-mediated mechanism that is very rapid as transcriptional activation occurs as early as 1hr post RUNX1 removal. This finding is consistent with previous results demonstrating that RUNX1 loss of function, either by shRNA knockdown or inhibition of the interaction between CBFβ and RUNX, has been shown to result in the emergence of a stem-like phenotype^7, 8, 10^. However, the immediacy and mechanism(s) underlying the acquisition of stemness and regulatory parameters of control cannot be directly addressed by conventional knockdown or approaches. Similarly, inhibition of the interaction between CBFβ and RUNX factors does not distinguish between the RUNX (RUNX1, RUNX2, RUNX3) family members and potential effects on other CBFβ functions.

**Figure 5.**
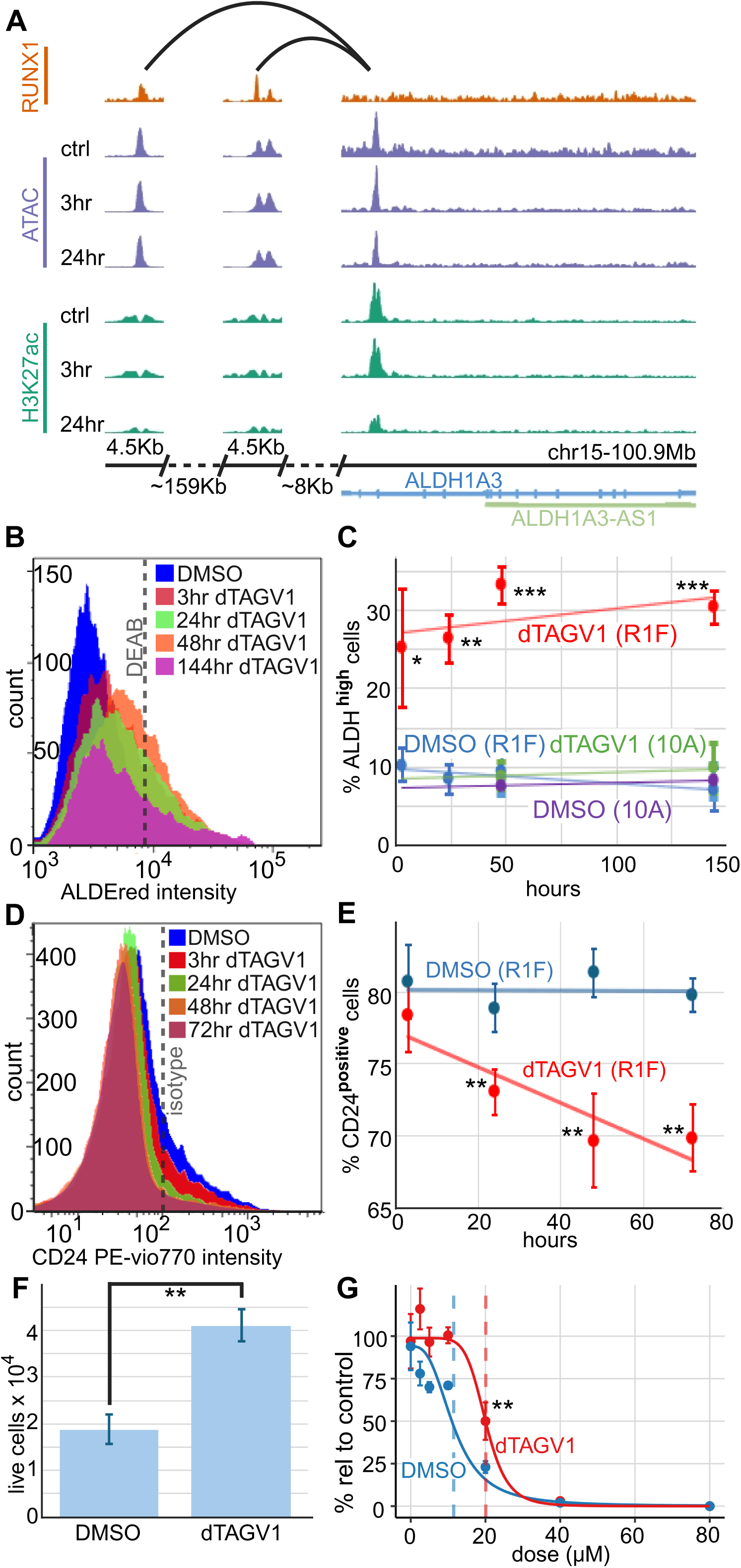
RUNX1 loss induces ALDH activity, cyclophosphamide resistance, and anchorage-independent survival. A) Chromosomal tracks depicting RUNX1 binding, chromatin contacts, ATAC and H3K27ac signals at chr15-101Mb region encompassing the ALDH1A3 locus that is upregulated upon the loss of RUNX1. B) Flow cytometry histograms of ALDH activity measured by an ALDEred assay at different timepoints following RUNX1 ablation. C) Quantification of ALDH high cells above DEAB-ALDH inhibitory controls indicate the percentage of cells that are ALDH bright in the DMSO baseline versus dTAGV1 treatment. D) Flow cytometry histograms of CD24 staining at different timepoints following RUNX1 ablation. E) Quantification of CD24+ cells above isotype controls indicates the percentage of CD24 high non-BCSCs in DMSO versus dTAGV1 treatment. F) MCF10A-R1F cells initially treated for 24hrs with DMSO or dTAGV1 were then subsequently placed in an anchorage-independent condition demonstrating an increased number of viable cells upon RUNX1 ablation. G) Relative MTS absorbance (450nm) as a measure of cell metabolism/ viability. Values are expressed as a percentage of vehicle control absorbance at multiple 4-hydroxycyclophosphamide (4-HC) dosages at 48hrs.

To assess the acquisition of stemlike properties, we utilized markers for BCSCs including ALDH activity and CD24 staining. ALDH activity is a primary marker for the generation and survival of BCSCs^24, 25^. As a negative control for ALDH activity in MCF10A-R1F cells, a DEAB inhibitor of ALDH1A1 and ALDH1A3 is used to gate ALDEred fluorescence signal (Figure 5B). Cells passing intensity threshold are considered ALDH^high^ BCSCs (AldeRed Assay, MilliporeSigma SCR150). Treatment of MCF10A-R1F cells with dTAGV1 establishes that acute loss of RUNX1 in mammary epithelial cells results in rapid emergence of ALDH^high^ BCSCs as early as 3hrs (Figure 5B,C). At 24hrs to 144hrs, there is a significant increase (p <0.01) in the number of ALDH^high^ BCSCs in dTAGV1 treated versus control. At 48hrs, there is an ∼3.4 fold increase in the number of ALDH^high^ BCSCs with no further increase in the number of ALDH^high^ BCSCs from 48 to 144hrs (Figure 5C). In contrast to ALDH activity, CD24 is a marker of the mammary epithelium and represents a negative indicator of BCSCs. CD24 was measured by flow cytometry on MCF10A-R1F treated with dTAGV1 or DMSO (Figure 5D). The percentage of CD24+ cells significantly (p<0.05) decreases to ∼69.6% in dTAGV1 treated cells at 48hrs and 72hrs, representing a ∼63% increase in CD24-BCSCs. Taken together, these findings demonstrate that RUNX1 loss diminishes epithelial identity and initiates a stem-like transformation program.

A prominent feature of BCSCs is their ability to survive in anchorage-independent conditions and resist alkylating chemotherapeutic agents. MCF10A-R1F pre-treated with dTAGV1 for 24hrs seeded in tumorsphere media in ultra-low attachment culture plates demonstrate increased cell viability of ∼2.1 fold (Figure 5F). Cyclophosphamide is frequently used for chemotherapy to treat breast cancer, and the drug’s effect is modified by ALDH activity. In MCF10A-R1F cells treated with 4-HC (4-hydroperoxycyclophosphamide, the active metabolite of the cyclophosphamide prodrug), cell viability is increased coincident with the loss of RUNX1 and subsequent increase in ALDH activity and/or positive cells. This chemoresistance was observed up to a dose of 20uM of 4-HC (Figure 5G). Together, these results suggest that loss of RUNX1 increases the number of BCSCs, enhances anchorage-independent survival, and promotes chemoresistance.

### RUNX1 loss regulates DNA damage responsiveness predominantly through promoters

Transcriptomic profiling revealed that RUNX1 depletion represses gene programs associated with DNA repair and checkpoint control. Functionally grouped ClueGO analysis of downregulated DEGs revealed significant enrichment for DNA damage response, cell cycle arrest, and genome stability pathways predominantly through promoters (Figure 4; Supplemental Figure 4). However, some DNA Damage Response (DDR) genes are putatively regulated through enhancer-promoter contacts. For example, FANCD2 is a central coordinator of DNA repair that responds to interstrand crosslinks and replication stress and is identified in our RUNX1 associated enhancer-promoter ABC analysis (Figure 6A). FANCD2 is contacted by one small (349bp) CRE located ∼11kb from the gene TSS (Figure 6A). This RUNX1-bound CRE is associated with an enhancer defined by broader ATAC and H3K27ac signals that interact with the proximal promoter of FANCD2. Because the promoter itself lacks RUNX1 occupancy, these results suggest that the presence of RUNX1 at the enhancer facilitates the activation of gene expression.

**Figure 6.**
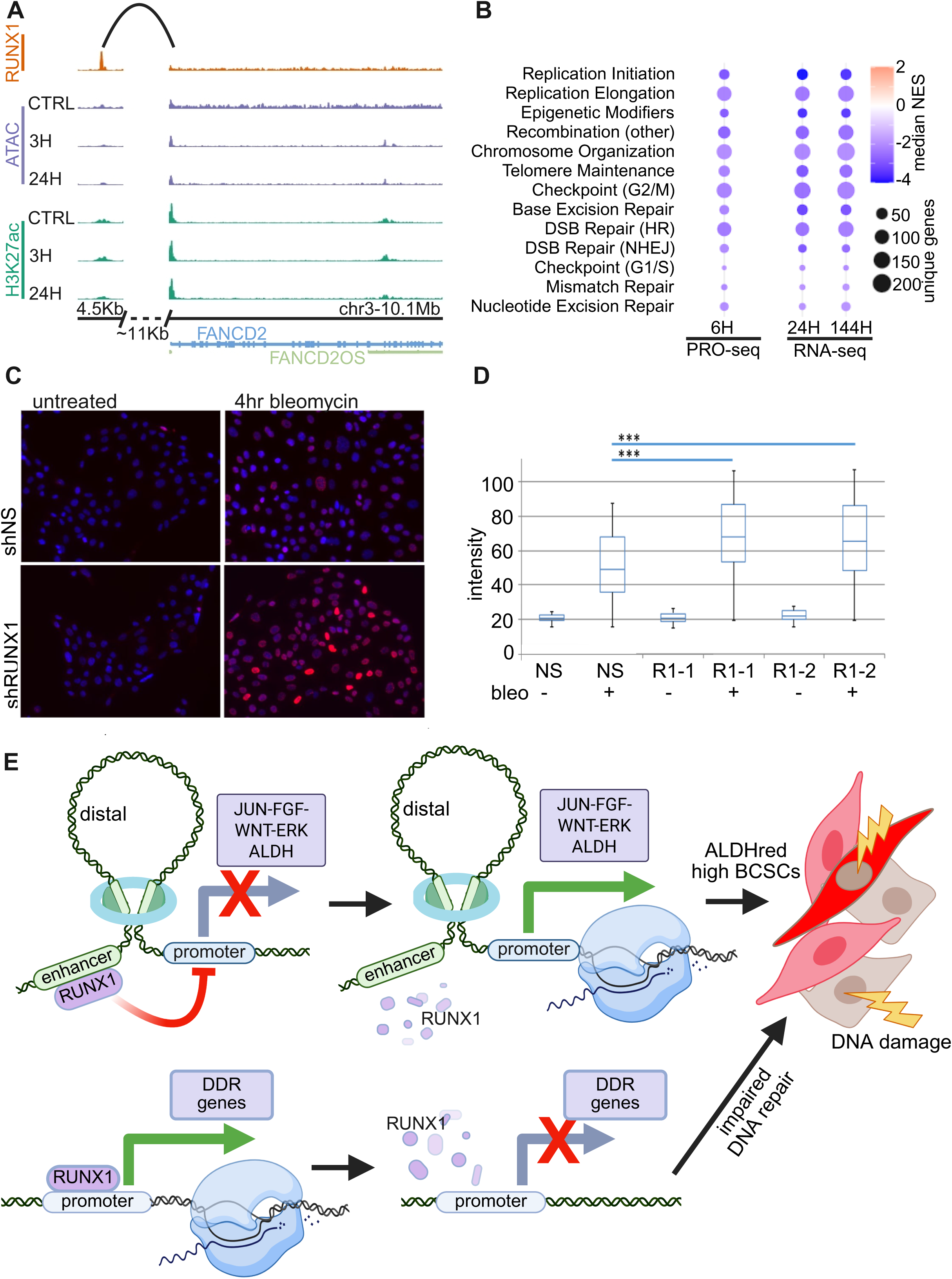
RUNX1 ablation results in altered expression of DNA damage response genes and increases yH2AX accumulation. A) Chromosomal tracks depicting RUNX1 binding, chromatin contacts, ATAC and H3K27ac signals at chr3-10Mb region encompassing the FANCD2 locus that is downregulated upon the loss of RUNX1. B) Gene Set Enrichment Analysis demonstrates that RUNX1 ablation results in negative enrichment scores for DNA damage response (DDR) related pathways. Pathways belonging to categories of DDR are term-reduced as shown in Supplementary Table 1. C) Immunofluorescence demonstrating yH2AX accumulation in RUNX1 deficient MCF10A treated with bleomycin for 4hrs versus control (untreated). D) Quantification of yH2AX signal intensity per nucleus in RUNX1 deficient cells with or without bleomycin treatment. Non-silencing shRNA (NS)-0hr bleomycin, n=2500; NS-4hr bleomycin, n=1465; shRUNX1-1 0hr bleomycin, n=1659; shRUNX1-construct1 (shRUNX1-1), 4hr bleomycin, n=2355; shRUNX1-construct2 (shRUNX1-2), 0hr bleomycin, n=1692; shRUNX1-construct2 (shRUNX1-2), 4hr bleomycin, n=1391; Statistical significance was determined by t-test, *** p <0.001. E) Schematic depiction of consequences of RUNX1 removal at distal enhancer- and promoter-related gene expression.

Following RUNX1 depletion, transcriptional downregulation of *FANCD2* and many other repair genes was evident as early as 6 hours in PRO-seq data and became apparent in RNA-seq by 24 hours (Figure 6B, 6C). Consistent with impaired DNA repair, γH2AX accumulation, a marker of unrepaired DNA damage^26^ increased progressively in RUNX1-depleted cells. Immunofluorescence after bleomycin treatment, which induces DNA strand breaks, revealed reduced DNA repair competency compared to controls (Figure 6D). Beyond regulating DNA repair, RUNX1 also modulates oncogenic signaling networks. Previous analyses (Figures 4D–F) showed that genes in the FGF–JUN–ERK pathway are repressed under normal RUNX1 function and become transcriptionally activated upon RUNX1 loss, suggesting a repressive role at these loci. In contrast to its promoter-mediated activation of DNA repair genes, RUNX1 suppresses oncogenic pathways via enhancer-mediated regulatory mechanisms. Upon RUNX1 loss, these oncogenic programs are derepressed, while genome maintenance systems are downregulated. Together, these findings indicate that RUNX1 coordinates enhancer–promoter networks that safeguard epithelial integrity and prevent early transformation (Figure 6E).

Defective repair response due to RUNX1 degradation is indicated by decreased expression of critical genes known to be involved in DNA repair and pathway inactivation as determined by GSEA of the DIG & DEG subset (Figure 6B, Supplementary Table 1). The alteration in DNA damage response pathways is transcriptionally evident at 6hrs in PRO-seq analyses, but not evident as steady-state RNA levels in RNA-seq analyses until 24hrs. An increase in γH2AX, is associated with a progressive accumulation of DNA damage^26^. In RUNX1 depleted MCF10A cells immunofluorescence staining relative to control exhibited impaired competency to repair DNA damage upon treatment with bleomycin, a chemotherapeutic agent that induces single and double strand breaks (Figure 6C, D). Removal of RUNX1 rapidly eliminates repression of genes in the FGF-JUN-ERK pathway potentially via enhancer-promoter interactions. In contrast, RUNX1 promoter-driven activation of DNA damage response is impaired upon its removal. These results strongly suggest that the loss of RUNX1 protein in normal breast cells is a precipitating event leading to widespread gene expression changes resulting in the loss of normal epithelial phenotype and acquisition of traits associated with tumor initiation and cancer progression (Figure 6E).

## Discussion

Our findings demonstrate an obligatory requirement for the RUNX1 transcription factor in breast cancer tumor suppression. Conventional knockdown strategies (e.g., siRNA and shRNA approaches) require extensive time for transcript and protein turnover resulting in ambiguities to define direct cancer-compromised regulatory mechanisms. In contrast, the degron-mediated strategy reported here immediately, selectively, and completely eliminates RUNX1, enabling temporal resolution of direct transcriptional consequences. There is an acute loss of normal mammary epithelial gene transcriptional initiation as early as 1hr after RUNX1 ablation, followed by an eventual acquisition of mesenchymal phenotypic properties. The approach defines a core cohort of RUNX1-responsive genes that are required to epigenetically regulate biological control in mammary epithelial cells (Supplementary Table 1). Our results suggest that RUNX1 drives promoter and distal enhancer-mediated gene expression programs supporting key pathways involved in sustaining the normal mammary epithelial phenotype. We have identified epigenetic remodeling of the chromatin landscape (histone modifications and higher-order chromatin organization (reviewed in^27, 28, 29^) at RUNX1 associated sites that modify the expression of genes involved in DNA damage responsiveness, plasticity, and stemness. The resulting transient or sustained change in transcriptional networks could be a key event in loss of epithelial phenotype and the induction of oncogenesis. Taken together, these results suggest that RUNX1 enhancer-promoter driven mechanisms mediate key hallmarks of cancer.

### RUNX1 distal enhancers regulate suppression of EMT and ALDH-active breast cancer stemness

Epithelial to Mesenchymal Transition (EMT) or partial EMT occurs during breast cancer initiation with modified cellular architecture and morphology as well as increased cell motility, cell survival, stemness, tumor initiating capacity, and drug resistance^30, 31, 32^. Residence of tumor cells at metastatic sites is frequently associated with a Mesenchymal to Epithelial Transition^33^. We find that the initial event upon the removal of RUNX1 is the rapid alteration of promoter and distal enhancer-mediated control of genes critical for EMT and pro-stemness signaling (e.g., FGF2^34, 35, 36^, DMBT1^37, 38, 39^, ALDH1A3^23^, and FANCD2^40^) (Figures 4 and 5). This analysis reveals activation of the TGF-β (transforming growth factor beta), FGF (fibroblast growth factor), MAPK (Mitogen Activated Protein Kinase), Wnt (Wingless/Int-1), NFkB (Nuclear Factor kappa B) pathways immediately following RUNX1 loss. These pathways are well-known inducers of EMT^41, 42, 43, 44, 45, 46^, and are altered upon the acute loss of RUNX1 in the absence of exogenous cytokines.

Consistent with the phenotype determining role of RUNX1 in multiple cell types^47, 48^ disrupting normal RUNX1 functioning in the AML-ETO translocation fusion protein (also known as RUNX1-RUNX1T1) impairs myeloid lineage determination^14^. Stengel et al. demonstrated that upon degradation of the AML1-ETO oncoprotein in leukemia cells, RUNX1 re-occupies a core set of genomic sites that are functionally linked to the myeloid lineage ^49^. In contrast, in normal mammary epithelial cells RUNX1 suppresses tumorigenesis and is the predominant RUNX factor with a minimal amount of RUNX2 or RUNX3 to compete for functional target gene regulation. Upon the degradation of AML-ETO in leukemia, there was a minimal change in global H3K27ac signal except at genomic sites where RUNX1 reoccupied a core set of genes. However, RUNX1 degradation in normal mammary epithelial cells results in acute alterations in H3K27ac and/or chromatin accessibility (ATAC-seq) at distal enhancers. Upon degradation of AML1-ETO in the cancer-compromised leukemia context there is a predominant activation at sites of RUNX1 reoccupancy^49^. Conversely, we find that RUNX1 loss in the normal mammary epithelial phenotype results in transcriptional repression. The tendency towards downregulation of transcriptional initiation and steady-state expression at early timepoints is functionally associated with a reduction in chromatin accessibility (ATAC-seq) and H3K27ac fold enrichment at regulatory sites. While RUNX1 is present at enhancers and promoters, this rapid chromatin alteration is primarily observed at enhancers consistent with engagement of Higher-order Chromatin Organization (HCO) in transcriptional control.

### Loss of RUNX1 results in metabolic reprogramming consistent with stemness and drug resistance

Reinforcing RUNX1 as a tumor suppressor, one of the earliest events after RUNX1 loss is the significant upregulation of several ALDH isoforms, indicating a shift in cellular metabolism favoring CSC survival (Figure 4 and 5). Elevated ALDH enzyme activity promotes detoxification of reactive lipid aldehydes, protects cells from DNA damage, and enhances survival under stress conditions including oxidative stress and chemotherapy^24, 50, 51^. Gene expression programs linked to these processes are identified to be rapidly transcriptionally initiated upon RUNX1 ablation and are subsequently reflected at the steady-state mRNA level (Figure 2). Clinical relevance for the activation of these stemness-related pathways upon the loss of RUNX1 is consistent with increased chemotherapy resistance against therapeutic agents including cyclophosphamide (Figure 5G). Heightened ALDH activity has been shown to neutralize toxic byproducts of lipid peroxidation, thereby avoiding ferroptosis or other forms of oxidative cell death^52, 53, 54^. Lipid metabolism and oxidative responsiveness are among the initial dominant pathways altered upon RUNX1 loss (Figure 4, Supplementary figure 4). This metabolic resilience, together with the activation of partial-EMT, contributes to the BCSC-like state of RUNX1-deficient cells and provides competency to withstand environmental (e.g. anchorage-independent survival) and therapeutic challenges (e.g. chemotherapy).

### RUNX1 mediates DNA damage responsiveness through promoters

RUNX1 functions as a tumor suppressor in several cell types including mammary epithelial cells. While disruption of RUNX1 has been shown to promote DNA Damage Response (DDR) competency particularly in leukemia^55, 56, 57, 58^, its role in breast cancer DDR is unclear. RUNX1 is activated in response to DNA damage and interacts with the ATM/ATR–Chk1/Chk2 signaling pathways^55^. In this study, we have identified that the RUNX1 mediated promoter and enhancer activities are key mechanisms regulating these pathways in the absence of exogenous DNA damage agents. Similarly, in leukemia the aberrant expression of AML1-ETO results in the suppression of DNA damage repair pathways and a corresponding increase in γH2AX foci, a marker of DNA damage^59^. We determine that the loss of RUNX1 in normal mammary epithelial cells leads to transcriptional repression of DDR genes observed as early as 6 hours after RUNX1 removal, and a marked reduction in steady-state expression levels by 24 hours potentially increasing susceptibility to secondary oncogenic mutations as suggested by an increased accumulation of γH2AX. These findings are consistent with induction of breast cancer-related stemness due to the loss of BRCA1^60^. Attenuated DNA damage response promotes the survival of cancer stem cells^61^. The ability of cancer stem cells to tolerate or exploit this genomic instability may further promote tumor progression^62^.

### RUNX1-enhancer and promoter mediated epigenetic tumor suppression

Evidence from these studies utilizing the power of degron-mediated RUNX1 ablation supports RUNX1 as an epigenetic tumor suppressor. Epigenetic control of RUNX1 tumor suppression includes immediate, transient, progressive and sustained parameters that include chromatin states (ChIP-seq) and chromatin organization reflected by openness (ATAC-seq) and higher order structure (Micro-C). Transcriptional initiation (PRO-seq) and steady-state gene expression (RNA-seq) reflect epigenetically mediated consequences for RUNX1 engagement in tumor suppression. Modified cell morphology, metabolic control, increased breast cancer stemness, anchorage-independent survival, chemotherapy resistance, and perturbed DNA damage responsiveness are observed upon the loss of RUNX1. The overarching consequence of RUNX1 loss is the acquisition of phenotypic characteristics associated with the hallmarks of cancer and loss of mammary epithelial properties. While support for RUNX1-mediated tumor suppression in mammary epithelial cells is compelling, there is a requirement for further interrogation of specific regulatory mechanisms that are operative in these transitions, and RUNX1 tumor promoter activity that has been reported in late-stage triple negative breast tumors^63^. It will be informative to expand understanding of epigenetic parameters of RUNX1 tumor suppression that include cell cycle control, mitotic gene bookmarking, higher-order chromatin organization, and DNA methylation.

## Materials and Methods

### Cell culture

MCF-10A^64^ cells were purchased from ATCC and grown in DMEM:F12 (Hyclone: SH30271, Thermo Fisher Scientific, Waltham, MA, USA) with 5% v/v horse serum (Gibco: 16050, Thermo Fisher Scientific) + 10 μg/ml human insulin (Sigma Aldrich, St. Louis, MO, USA: I-1882) + 20 ng/ml recombinant hEGF (Peprotech, Rocky Hill, NJ, USA: AF-100-15) + 100 ng/ml cholera toxin (Sigma Aldrich: C-8052) + 0.5 μg/ml hydrocortisone (Sigma Aldrich: H-0888) + 50 IU/ml penicillin/ 50 μg/ml streptomycin and 2 mM glutamine (Life Technologies, Carlsbad, CA, USA: 15140-122 and 25030-081, respectively). MCF-10CA1a cells were grown in DMEM:F12 with 5% v/v horse serum, 50 IU/ml penicillin/ 50 μg/ml streptomycin, and 2 mM glutamine.

### CRISPR-mediated knockin of FKBP12 onto RUNX1

To directly study the initial, progressive, and sustained consequences of the loss of the RUNX1 protein at specific timepoints, we employed a CRISPR-based strategy to tag the endogenous RUNX1 gene with an HA-HA-FKBP12^F36V^-P2A-mCherry sequence at the C-terminus. The P2A sequence is a self-cleaving domain that separates the RUNX1-HA-HA-FKBP12^F36V^ from the mCherry. Briefly, crRNA targeting the 3′ end of *RUNX1* was annealed with tracrRNA (IDT). Annealed gRNAs were assembled into RNP complexes by incubation with Alt-R S.p. Cas9 Nuclease V3 (IDT) at room temperature. RNP complexes and equimolar HDR plasmid targeting the 3′ exon of *RUNX1* and containing *FKBP12^F36V^-2xHA-P2A-mCherry* were electroporated into MCF10A cells. Successfully edited MCF10A-RUNX1-linker-HA-HA-FKBP12^F36V^-mCherry (MCF10A-R1F) cells were isolated by fluorescence-activated cell sorting using the co-expressed mCherry and screened for homologous recombination by PCR, western blot analysis, and immunofluorescence imaging. The FKBP12^F36V^ epitope is ubiquitinated upon the addition of a small molecule designated dTAGV1 that recruits the Von Hippel-Lindau E3-ligase complex (Figure 1A).

### RNA-seq analysis

Total RNA was isolated from cells using Trizol (Life Technologies) and purified using the Direct-zol RNA kit (Zymo Research, Irvine, CA, USA: R2050) according to the manufacturer’s instructions. RNA quantity was assessed using a Nanodrop2000 (Thermo Scientific, Lafayette, CO) and Qubit HS RNA assay (Thermo Fisher Scientific), and RNA 6000 Nano Kit with the Agilent 2100 Bioanalyzer (Agilent Technologies, Santa Clara, CA). Total RNA was depleted of ribosomal RNA, reverse transcribed and strand-specific adapters added following manufacturer’s protocol (SMARTer Stranded Total RNA Sample Prep Kit - HI Mammalian, Catalog number: 634876). Generated cDNA libraries were assayed for quality using the High Sensitivity DNA Kit on the Agilent 2100 Bioanalyzer (Agilent Technologies) then sequenced as paired-end 150 bp reads (Novaseq6000, Novogene).

RNA-seq experiments were performed in triplicate. Raw paired-end RNA-seq reads were processed using the nf-core/rnaseq pipeline with default parameters^65^. Reads were trimmed using Trim Galore and aligned to the human genome (hg38) using STAR, followed by transcript quantification with Salmon. Differential gene expression analysis was performed using DESeq2. Genes were considered significantly differentially expressed if they exhibited an adjusted *p*-value (*padj*) < 0.05 and an absolute fold-change > 1.5. RNA-Seq datasets have been deposited in the Gene Expression Omnibus (GEO) under accession code GSE314305.

### PROseq analysis

PRO-seq experiments were performed as previously reported^49^. Adapter trimming was performed with Trim Galore, and paired-end reads were aligned to the hg38 genome using Bowtie2. Following alignment, the R1 mate was extracted to represent the engaged RNA polymerase position. Strand-specific bedGraph files were generated and read counts were quantified using featureCounts across defined genomic regions: promoter-proximal regions (TSS ±250 bp) and gene bodies (TSS - TES, extended by 250 bp on each side). Differential expression analysis was performed on the gene body counts using DESeq2. Genes were considered significantly differentially initiated if they exhibited adjusted *p*-value (*padj*) < 0.05 and an absolute fold-change > 1.5. PRO-Seq datasets have been deposited in the Gene Expression Omnibus (GEO) under accession code GSE314312.

### ATAC-seq analysis

ATAC-seq was performed according to the OMNI-ATACseq protocol as previously described^66^ with an increased nuclear isolation time of 5 minutes instead of 3 minutes. ATAC-seq preprocessing was performed using custom scripts implemented according to the ENCODE ATAC-seq pipeline guidelines, with minor modifications to accommodate our experimental design^67^. FASTQ files were downsampled to match the lowest depth across samples (∼50M reads). Paired-end reads were aligned to the hg38 genome with Bowtie2, and alignments were filtered using SAMtools to remove mitochondrial reads, PCR duplicates, and low-quality alignments. Filtered BAM files were converted to TAGAlign format, and Tn5 insertion sites were adjusted to account for transposase offset (+4 bp for the positive strand and –5 bp for the negative strand). Biological replicates were pooled, and peaks were called using MACS2, following Encode standard parameters. Differential accessibility analysis between conditions was performed using DiffBind (FDR < 0.05)^68^. ATAC-Seq datasets have been deposited in the Gene Expression Omnibus (GEO) under accession code GSE314301.

### ChIP-seq analysis

ChIP-seq was performed as previously described^69^. We performed independent replicates for DMSO and dTAGV-1 treated MCF10A-RUNX1-FKBP cells for H3K27ac (Abcam, ab4729). These ChIP-seq datasets have been deposited in the Gene Expressions Omnibus (GEO) under accession code. H3K27ac ChIP-seq data was processed using the nf-core/chipseq pipeline^65^ with default parameters. Reads were aligned to hg38, followed by filtering and duplicate removal. Peaks were called relative to matched input controls using MACS2, as implemented within the pipeline. Differential enrichment analysis of H3K27ac peaks between conditions was performed using DiffBind^68^ with default normalization and contrast settings. ChIP-Seq datasets have been deposited in the Gene Expression Omnibus (GEO) under accession code GSE314307.

### Enhancer Definitions

Candidate enhancers were defined by intersecting multiple regulatory annotations. First, ChromHMM-annotated enhancer states in MCF10A cells were overlapped with ENCODE candidate cis-regulatory elements (cCREs) classified as distal enhancers^67^. This intersection yielded a high-confidence set of putative enhancer regions. To focus on direct targets of RUNX1, regions overlapping RUNX1 ChIP-seq peaks were extracted. These RUNX1-associated enhancers were then linked to target promoters using MCF10A-specific Activity-By-Contact (ABC) model predictions^70^, retaining enhancer–gene pairs with an ABC score > 0.02.

### ClueGO Pathway Enrichment Analysis

To investigate the dynamic biological consequences of RUNX1 degradation, we performed pathway enrichment analysis using the ClueGO plugin (v2.5.10) within Cytoscape (v3.10.3). Enrichment was conducted across Gene Ontology (GO) Biological Processes, KEGG pathways, and Reactome databases, enabling the mapping of temporal shifts in transcriptional programs associated with RUNX1 loss. All databases were current as of April 16, 2025.

### Coimmunoprecipitation

For endogenous co-immunoprecipitation, cells were plated in 10-cm dishes and lysed in RIPA buffer supplemented with protease inhibitors and MG132. Lysates were sonicated and clarified by centrifugation at 10,000 rpm, and protein concentration was determined by BCA assay. For each immunoprecipitation, 1 mg of total protein was incubated overnight at 4 °C with either HA agarose beads or control agarose beads. Beads were then washed four times with NETN buffer, and bound proteins were eluted by boiling in Laemmli sample buffer. Immunoprecipitates were analyzed by SDS PAGE and immunoblotting using RUNX1 (CST, 4336), CBFβ (Abcam, ab231345), and TLE1 (EMD Millipore, ABD119) antibodies, input lysates were probed with an HA antibody.

### Western Blot analysis

Cells were lysed in RIPA buffer supplemented with protease 231inhibitors and MG132 and sonicated to shear DNA. Cleared lysates were resolved by SDS-PAGE, transferred to PVDF membrane and probed with specified antibodies. For RUNX2, CBFβ, and CTCF westerns the OMNI-ATAC protocol was used prior to lysing in RIPA. Antibodies: *α-*HA (Abcam ab18181), α-RUNX1- (CST, 4336), α-CBFβ (Abcam, EPR6322), α-BACT (CST, 3700), α-RUNX2 (CST, D1L7F), α-CTCF (CST, 2899).

### Immunofluorescence/Immunohistochemistry

For Beta actin and F-actin staining, MCF10A-R1F were treated with DMSO or dTAGV1 for 144hrs and fixed in 4% formaldehyde for 10 minutes at room temperature, permeabilized with 0.25% triton for 10 minutes, blocked with 1% BSA for 1hr, stained overnight at 4oC with α-βACT (1:1000, Cell Signaling Technologies, 3700S), and then phalloidin-647 (ab176759, Abcam), secondary (1:2000, Invitrogen, A-11001), and DAPI (1:2000 Sigma, D9542) for 1hr at room temperature. For yH2AX analysis RUNX1 was knocked down as previously described and stained with yH2AX overnight at 4oC and counterstained with DAPI. Nuclei were segmented, and fluorescence intensity was measured using ImageJ.

### AldeRed Assay and Flow cytometry

Aldehyde dehydrogenase activity was determined using the AldeRed Detection Assay, Catalog number: SCR150. Briefly, MCF10A or MCF10A-RUNX1-FKBP12^F36V^ were treated with either DMSO or dTAGV-1 for the designated amount of time and then trypsinized and 1 million were incubated with ALDEred and split in half to incubate with or without the addition of DEAB inhibitor of ALDH activity. ALDH bright versus ALDH low was determined by using the DEAB controls for gating.

For CD24 staining, cells were washed and resuspended at a concentration of 1 million cells per 100 μL in PBS + 0.5% BSA. Cells were stained with CD24 (Miltenyi Biotec, 130-108-352) or isotype (Miltenyi Biotec, 130-124-062) antibodies at the manufacturer’s suggested concentration for 20 minutes at 4°C, washed and acquired using a MACSquant YVB.

### Anchorage-Independent Survival

MCF10A-R1F cells were treated with DMSO or dTAGV1 for 24hrs under standard growth conditions. These cells were accutased, PBS washed centrigued and resuspended in tumorsphere media (MammoCultTM Human Medium Kit, Stemcell Technologies, 05620) supplemented with heparin sulphate (Stemcell Technologies, 07980) and hydrocortisone (Stemcell Technologies, 74142) and 40uM mesh filtered to generate a single cell suspension. 2x10^3 Cells per 100ul were then plated in Nunclon Sphera ultralow attachment plates (Fisher Scientific, 12-566-430). After 24hrs under anchorage-independent conditions, resulting tumorspheres from 12 wells (from a 96 well plate) were combined and disaggregated using trypsin and counted using trypan blue exclusion.

### MTS assay

MCF10A-R1F cells were plated at 2x10^3 grown for 24hrs and then treated with DMSO or dTAGV1 for 24hrs, followed by 4-hydroperoxy cyclophosphamide (Cayman Chemical Company, 19527) at the specified concentrations for 48hrs. CCK8 (1:10, Sigma, 96992) was incubated for 1hr at 37oC and absorbance at 450nM was measured on a Biotek plate reader. The absorbance of surviving cells at each concentration was expressed as a percent of its corresponding vehicle DMSO control absorbance.

## Acknowledgements

We thank Genieve Brzoza, Joshua Mercurie, Adam Wineheimer, Stephen Foley, Lizzy Hahn, and Dorcas Lohese for valuable assistance with cell culture and feedback. Support for this work was provided by the National Cancer Institute (P01-CA240685 to GS, JS, and SF), the C. Perelman and A.J. Perelman endowments, the Intramural Research Program of the National Cancer Institute, Center for Cancer Research (ZIABC010309-24 to T.M), the Northern New England Clinical and Translational Research Network (GM115516 to GS), and the Vanderbilt-Ingram Cancer Center Genome Sciences Shared Resource (NCIP30CA68485, SH and KS). STR analysis and next-generation sequencing was in part done with the assistance of the Vermont Integrated Genomics Resource at the University of Vermont Cancer Center and Jordan Zhang, Misha Gattengo, and Sierra Wilson at Dovetail Genomics coordinated Micro-C library preparation and preliminary analysis. The contributions of the NIH author(s) were made as part of their official duties as NIH federal employees, are in compliance with agency policy requirements, and are considered Works of the United States Government. However, the findings and conclusions presented in this paper are those of the author(s) and do not necessarily reflect the views of the NIH or the U.S. Department of Health and Human Services.

## Author Contributions

Experimental Contributions provided by *Andrew J. Fritz, Haley Greenyer, Rahim Ullah, Louis Dillac, Priyanka Chavarkar, Abigail G. Person, Deli Hong, Kathleen Reed, Georgiy Zotkin, Sadie Korzec, Cong Gao, Miles Malik, Jonah Perelman, Alquassem Abuorquob, Jackson Del Porto*. Research design and data analysis provided by *Andrew J. Fritz, Haley Greenyer, Rahim Ullah, Louis Dillac, Deli Hong, Kathleen Reed, Georgiy Zotkin, Cong Gao, Martin Montecino, Jane B. Lian, Andre Van Wijnen Jessica Heath, Prachi Ghule, Seth Frietze, Kristy Stengel, Scott Hiebert, Thomas Misteli, Jonathan A. Gordon, Janet L. Stein, Gary S. Stein*. Writing of manuscript provided by *Andrew J. Fritz, Haley Greenyer*, *Emory Pacht, Rabail Toor, Kathleen Reed, Janine Warren, Georgiy Zotkin, Martin Montecino, Jane B. Lian, Andre Van Wijnen, Jessica Heath, Prachi Ghule, Seth Frietze, Kristy Stengel, Scott Hiebert, Thomas Misteli, Jonathan A. Gordon, Janet L. Stein, Gary S. Stein*

**Supplementary Figure 1. Degron-containing MCF10A-R1F robustly express key mammary epithelial genes.** The normalized counts in RNA-seq data is displayed for key genes in parental MCF10A and RUNX1-FKBPcontaining MCF10A-R1F.

**Supplementary Figure 2. Processed steady-state RNAs with higher intron/isoform density are expressed later, while differential transcriptional initiation is not coincident with intron/isoform density.** A) The absolute values of DEGs for up versus downregulated in PRO-seq and RNA-seq were compared and Wilcoxon rank-sum tests determined a stronger magnitude of transcriptional repression than upregulation. B) Partial spearman values standardizing for transcript length and genomic span (total length of the genomic locus including introns and exons) are displayed for genomic characteristics.

**Supplementary Figure 3. Differential peak calls for ATAC-seq and H3K27ac-seq across time and ChromHMM state emissions.** A) Volcano plots illustrate significant differential ATAC-seq peaks (FDR < 0.05) DMSO vs. dTAGV1 at 3, 24, and 144 hours. B) Volcano plots illustrate significant differential H3K27ac ChIP-seq peaks (FDR < 0.05) DMSO vs. dTAGV1 at 3, 24, and 144 hours. C) ChromHMM 20-state emission heatmap generated from indicated ChIP-seq and RNA-seq signals in MCF10A.

**Supplementary Figure 4. Functional organized network of biological terms of DIG/ DEGs upon RUNX1 loss.** A) Functionally grouped network ClueGo analysis for DIG/ DEG upon RUNX1 ablation. Color scale indicates padj values and bubble size indicates weighted percent association defined pathways. Heatmap of row normalized z-score values of select genes associated with B) FGF signaling and C) DNA damage response. Jaccard similarity matrix of D) upregulated and E) downregulated genes.

**Supplementary Figure 5. RUNX2 protein expression increases after 12hrs post RUNX1 ablation long after initial.** Western blot demonstrating that RUNX2 increases in protein expression but only after 12hrsof dTAGV1 treatment well after changes in gene expression. Exposure time for RUNX2 was significantly longer than RUNX1 indicating that RUNX2 expression is qualitatively less than RUNX1.

## REFERENCES

1. Ramaswamy S, Ross KN, Lander ES, Golub TR. A molecular signature of metastasis in primary solid tumors. Nat Genet 33, 49–54 (2003).

2. Hong D, et al. RUNX1-dependent mechanisms in biological control and dysregulation in cancer. J Cell Physiol 234, 8597–8609 (2019).

3. Hong D, et al. Runx1 stabilizes the mammary epithelial cell phenotype and prevents epithelial to mesenchymal transition. Oncotarget 8, 17610–17627 (2017).

4. Kadota M, et al. Delineating genetic alterations for tumor progression in the MCF10A series of breast cancer cell lines. PLoS One 5, e9201 (2010).

5. Banerji S, et al. Sequence analysis of mutations and translocations across breast cancer subtypes. Nature 486, 405–409 (2012).

6. Wang L, Brugge JS, Janes KA. Intersection of FOXO- and RUNX1-mediated gene expression programs in single breast epithelial cells during morphogenesis and tumor progression. Proc Natl Acad Sci U S A 108, E803–812 (2011).

7. Hong D, et al. Suppression of Breast Cancer Stem Cells and Tumor Growth by the RUNX1 Transcription Factor. Mol Cancer Res 16, 1952–1964 (2018).

8. Kulkarni M, et al. RUNX1 and RUNX3 protect against YAP-mediated EMT, stemness and shorter survival outcomes in breast cancer. Oncotarget 9, 14175–14192 (2018).

9. Sokol ES, Sanduja S, Jin DX, Miller DH, Mathis RA, Gupta PB. Perturbation-expression analysis identifies RUNX1 as a regulator of human mammary stem cell differentiation. PLoS Comput Biol 11, e1004161 (2015).

10. Fritz AJ, et al. RUNX1 and RUNX2 transcription factors function in opposing roles to regulate breast cancer stem cells. J Cell Physiol 235, 7261–7272 (2020).

11. Barutcu AR, et al. RUNX1 contributes to higher-order chromatin organization and gene regulation in breast cancer cells. Biochim Biophys Acta 1859, 1389–1397 (2016).

12. Hass MR, et al. Runx1 shapes the chromatin landscape via a cascade of direct and indirect targets. PLoS Genet 17, e1009574 (2021).

13. de la Serna IL, Ohkawa Y, Imbalzano AN. Chromatin remodelling in mammalian differentiation: lessons from ATP-dependent remodellers. Nat Rev Genet 7, 461–473 (2006).

14. Speck NA, Gilliland DG. Core-binding factors in haematopoiesis and leukaemia. Nat Rev Cancer 2, 502–513 (2002).

15. Rose JT, et al. Inhibition of the RUNX1-CBFbeta transcription factor complex compromises mammary epithelial cell identity: a phenotype potentially stabilized by mitotic gene bookmarking. Oncotarget 11, 2512–2530 (2020).

16. Yonezawa T, et al. The ubiquitin ligase STUB1 regulates stability and activity of RUNX1 and RUNX1-RUNX1T1. J Biol Chem 292, 12528–12541 (2017).

17. Debnath J, Muthuswamy SK, Brugge JS. Morphogenesis and oncogenesis of MCF-10A mammary epithelial acini grown in three-dimensional basement membrane cultures. Methods 30, 256–268 (2003).

18. Gross SM, et al. A multi-omic analysis of MCF10A cells provides a resource for integrative assessment of ligand-mediated molecular and phenotypic responses. Commun Biol 5, 1066 (2022).

19. Qu Y, et al. Evaluation of MCF10A as a Reliable Model for Normal Human Mammary Epithelial Cells. PLoS One 10, e0131285 (2015).

20. Gardel ML, Schneider IC, Aratyn-Schaus Y, Waterman CM. Mechanical integration of actin and adhesion dynamics in cell migration. Annu Rev Cell Dev Biol 26, 315–333 (2010).

21. van Bragt MP, Hu X, Xie Y, Li Z. RUNX1, a transcription factor mutated in breast cancer, controls the fate of ER-positive mammary luminal cells. Elife 3, e03881 (2014).

22. Vander Ark A, Cao J, Li X. TGF-beta receptors: In and beyond TGF-beta signaling. Cell Signal 52, 112–120 (2018).

23. McLean ME, et al. The Expanding Role of Cancer Stem Cell Marker ALDH1A3 in Cancer and Beyond. Cancers (Basel) 15, (2023).

24. Lavudi K, Nuguri SM, Pandey P, Kokkanti RR, Wang QE. ALDH and cancer stem cells: Pathways, challenges, and future directions in targeted therapy. Life Sci 356, 123033 (2024).

25. Charafe-Jauffret E, et al. Breast cancer cell lines contain functional cancer stem cells with metastatic capacity and a distinct molecular signature. Cancer Res 69, 1302–1313 (2009).

26. Rogakou EP, Pilch DR, Orr AH, Ivanova VS, Bonner WM. DNA double-stranded breaks induce histone H2AX phosphorylation on serine 139. J Biol Chem 273, 5858–5868 (1998).

27. Berezney R. Regulating the mammalian genome: the role of nuclear architecture. Adv Enzyme Regul 42, 39–52 (2002).

28. Yang JH, Hansen AS. Enhancer selectivity in space and time: from enhancer-promoter interactions to promoter activation. Nat Rev Mol Cell Biol 25, 574–591 (2024).

29. Robson MI, Ringel AR, Mundlos S. Regulatory Landscaping: How Enhancer-Promoter Communication Is Sculpted in 3D. Mol Cell 74, 1110–1122 (2019).

30. Kalluri R, Weinberg RA. The basics of epithelial-mesenchymal transition. J Clin Invest 119, 1420–1428 (2009).

31. Allgayer H, et al. Epithelial-to-mesenchymal transition (EMT) and cancer metastasis: the status quo of methods and experimental models 2025. Mol Cancer 24, 167 (2025).

32. Huang Y, Hong W, Wei X. The molecular mechanisms and therapeutic strategies of EMT in tumor progression and metastasis. J Hematol Oncol 15, 129 (2022).

33. Ribatti D, Tamma R, Annese T. Epithelial-Mesenchymal Transition in Cancer: A Historical Overview. Transl Oncol 13, 100773 (2020).

34. Maehara O, et al. Fibroblast growth factor-2-mediated FGFR/Erk signaling supports maintenance of cancer stem-like cells in esophageal squamous cell carcinoma. Carcinogenesis 38, 1073–1083 (2017).

35. Zhao Q, et al. FGFR inhibitor, AZD4547, impedes the stemness of mammary epithelial cells in the premalignant tissues of MMTV-ErbB2 transgenic mice. Sci Rep 7, 11306 (2017).

36. Bhat V, Piaseczny M, Goodale D, Patel U, Sadri A, Allan AL. Lung-derived soluble factors support stemness/plasticity and metastatic behaviour of breast cancer cells via the FGF2-DACH1 axis. Clin Exp Metastasis 41, 717–731 (2024).

37. Zhang CX. The protective role of DMBT1 in cervical squamous cell carcinoma. Kaohsiung J Med Sci 35, 739–749 (2019).

38. Singh P, et al. Squamous cell carcinoma subverts adjacent histologically normal epithelium to promote lateral invasion. J Exp Med 218, (2021).

39. Braidotti P, et al. DMBT1 expression is down-regulated in breast cancer. BMC Cancer 4, 46 (2004).

40. Wang H, et al. BRCA1/FANCD2/BRG1-Driven DNA Repair Stabilizes the Differentiation State of Human Mammary Epithelial Cells. Mol Cell 63, 277–292 (2016).

41. Zhang J, et al. TGF-beta-induced epithelial-to-mesenchymal transition proceeds through stepwise activation of multiple feedback loops. Sci Signal 7, ra91 (2014).

42. Brown WS, Tan L, Smith A, Gray NS, Wendt MK. Covalent Targeting of Fibroblast Growth Factor Receptor Inhibits Metastatic Breast Cancer. Mol Cancer Ther 15, 2096–2106 (2016).

43. Shin S, Dimitri CA, Yoon SO, Dowdle W, Blenis J. ERK2 but not ERK1 induces epithelial-to-mesenchymal transformation via DEF motif-dependent signaling events. Mol Cell 38, 114–127 (2010).

44. Xue W, Yang L, Chen C, Ashrafizadeh M, Tian Y, Sun R. Wnt/beta-catenin-driven EMT regulation in human cancers. Cell Mol Life Sci 81, 79 (2024).

45. Dongre A, Weinberg RA. New insights into the mechanisms of epithelial-mesenchymal transition and implications for cancer. Nat Rev Mol Cell Biol 20, 69–84 (2019).

46. Oh A, et al. NF-kappaB signaling in neoplastic transition from epithelial to mesenchymal phenotype. Cell Commun Signal 21, 291 (2023).

47. Mevel R, Draper JE, Lie ALM, Kouskoff V, Lacaud G. RUNX transcription factors: orchestrators of development. Development 146, (2019).

48. Rozen EJ, Ozeroff CD, Allen MA. RUN(X) out of blood: emerging RUNX1 functions beyond hematopoiesis and links to Down syndrome. Hum Genomics 17, 83 (2023).

49. Stengel KR, Ellis JD, Spielman CL, Bomber ML, Hiebert SW. Definition of a small core transcriptional circuit regulated by AML1-ETO. Mol Cell 81, 530–545 e535 (2021).

50. Grimley E, et al. Aldehyde dehydrogenase inhibitors promote DNA damage in ovarian cancer and synergize with ATM/ATR inhibitors. Theranostics 11, 3540–3551 (2021).

51. Poturnajova M, Kozovska Z, Matuskova M. Aldehyde dehydrogenase 1A1 and 1A3 isoforms - mechanism of activation and regulation in cancer. Cell Signal 87, 110120 (2021).

52. Wu Y, Franzmeier S, Liesche-Starnecker F, Schlegel J. Enhanced Sensitivity to ALDH1A3-Dependent Ferroptosis in TMZ-Resistant Glioblastoma Cells. Cells 12, (2023).

53. Wu W, et al. Lipid Peroxidation Plays an Important Role in Chemotherapeutic Effects of Temozolomide and the Development of Therapy Resistance in Human Glioblastoma. Transl Oncol 13, 100748 (2020).

54. Singh S, et al. Aldehyde dehydrogenases in cellular responses to oxidative/electrophilic stress. Free Radic Biol Med 56, 89–101 (2013).

55. Samarakkody AS, Shin NY, Cantor AB. Role of RUNX Family Transcription Factors in DNA Damage Response. Mol Cells 43, 99–106 (2020).

56. Wu D, Ozaki T, Yoshihara Y, Kubo N, Nakagawara A. Runt-related transcription factor 1 (RUNX1) stimulates tumor suppressor p53 protein in response to DNA damage through complex formation and acetylation. J Biol Chem 288, 1353–1364 (2013).

57. Satoh Y, et al. C-terminal mutation of RUNX1 attenuates the DNA-damage repair response in hematopoietic stem cells. Leukemia 26, 303–311 (2012).

58. Wang CQ, et al. Disruption of Runx1 and Runx3 leads to bone marrow failure and leukemia predisposition due to transcriptional and DNA repair defects. Cell Rep 8, 767–782 (2014).

59. Krejci O, et al. p53 signaling in response to increased DNA damage sensitizes AML1-ETO cells to stress-induced death. Blood 111, 2190–2199 (2008).

60. Kim H, Lin Q, Yun Z. BRCA1 regulates the cancer stem cell fate of breast cancer cells in the context of hypoxia and histone deacetylase inhibitors. Sci Rep 9, 9702 (2019).

61. Chen Y, et al. Quiescence and attenuated DNA damage response promote survival of esophageal cancer stem cells. J Cell Biochem 113, 3643–3652 (2012).

62. Vitale I, Manic G, De Maria R, Kroemer G, Galluzzi L. DNA Damage in Stem Cells. Mol Cell 66, 306–319 (2017).

63. Khan AS, Campbell KJ, Cameron ER, Blyth K. The RUNX/CBFbeta Complex in Breast Cancer: A Conundrum of Context. Cells 12, (2023).

64. Soule HD, et al. Isolation and characterization of a spontaneously immortalized human breast epithelial cell line, MCF-10. Cancer Res 50, 6075–6086 (1990).

65. Ewels PA, et al. The nf-core framework for community-curated bioinformatics pipelines. Nat Biotechnol 38, 276–278 (2020).

66. Corces MR, et al. An improved ATAC-seq protocol reduces background and enables interrogation of frozen tissues. Nat Methods 14, 959–962 (2017).

67. Consortium EP. An integrated encyclopedia of DNA elements in the human genome. Nature 489, 57–74 (2012).

68. Ross-Innes CS, et al. Differential oestrogen receptor binding is associated with clinical outcome in breast cancer. Nature 481, 389–393 (2012).

69. O’Geen H, Frietze S, Farnham PJ. Using ChIP-seq technology to identify targets of zinc finger transcription factors. Methods Mol Biol 649, 437–455 (2010).

70. Reed KSM, et al. Genome reorganization and its functional impact during breast cancer progression. bioRxiv, (2025).

