## Supplementary Figures for "Acute degron-mediated RUNX1 loss reprograms enhancer activity to epigenetically drive epithelial destabilization and initiate cancer hallmarks"

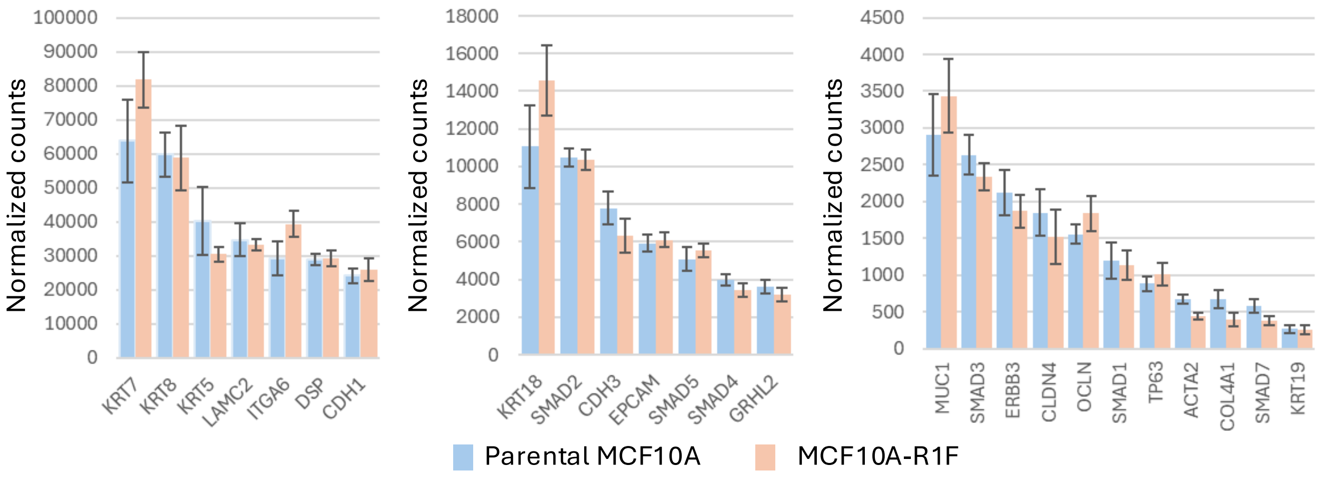


**Figure S1. Degron-containing MCF10A-R1F robustly express key mammary epithelial genes.** The normalized counts in RNAseq data is displayed for key genes in parental MCF10A and RUNX1-FKBP containing MCF10A-R1F.

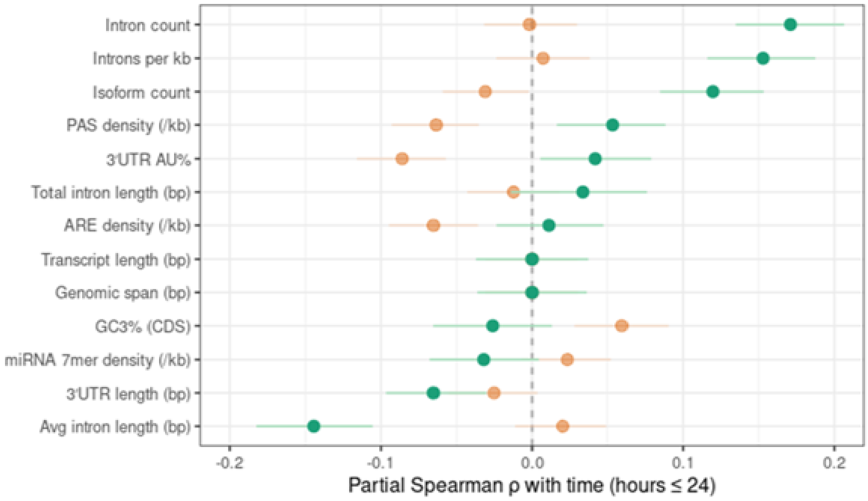


**Figure S2. Processed steady-state RNAs with higher intron/isoform density are expressed later, while differential transcriptional initiation is not coincident with intron/isoform density.** Partial spearman values standardizing for transcript length and genomic span (total length of the genomic locus including introns and exons) are displayed for genomic characteristics.

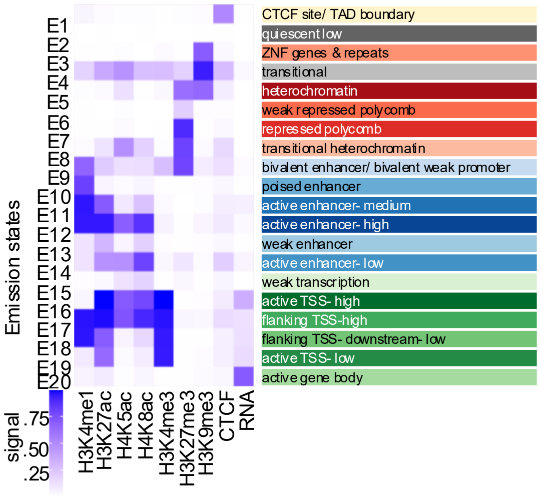

E1
E2
E3
E4
E5
E6
E7
E8
E9
E10
E11
E12
E13
E14
E15
E16
E17
E18
E19
E20
Emission states
signal
.75
.50
.25
H3K4me1
H3K27ac
H4K5ac
H4K8ac
H3K4me3
H3K27me3
H3K9me3
CTCF
RNA
CTCF site/ TAD boundary
quiescent low
ZNF genes & repeats
transitional
heterochromatin
weak repressed polycomb
repressed polycomb
transitional heterochromatin
bivalent enhancer/ bivalent weak promoter
poised enhancer
active enhancer- medium
active enhancer- high
weak enhancer
active enhancer- low
weak transcription
active TSS- high
flanking TSS-high
flanking TSS- downstream- low
active TSS- low
active gene body


**Figure S3.**

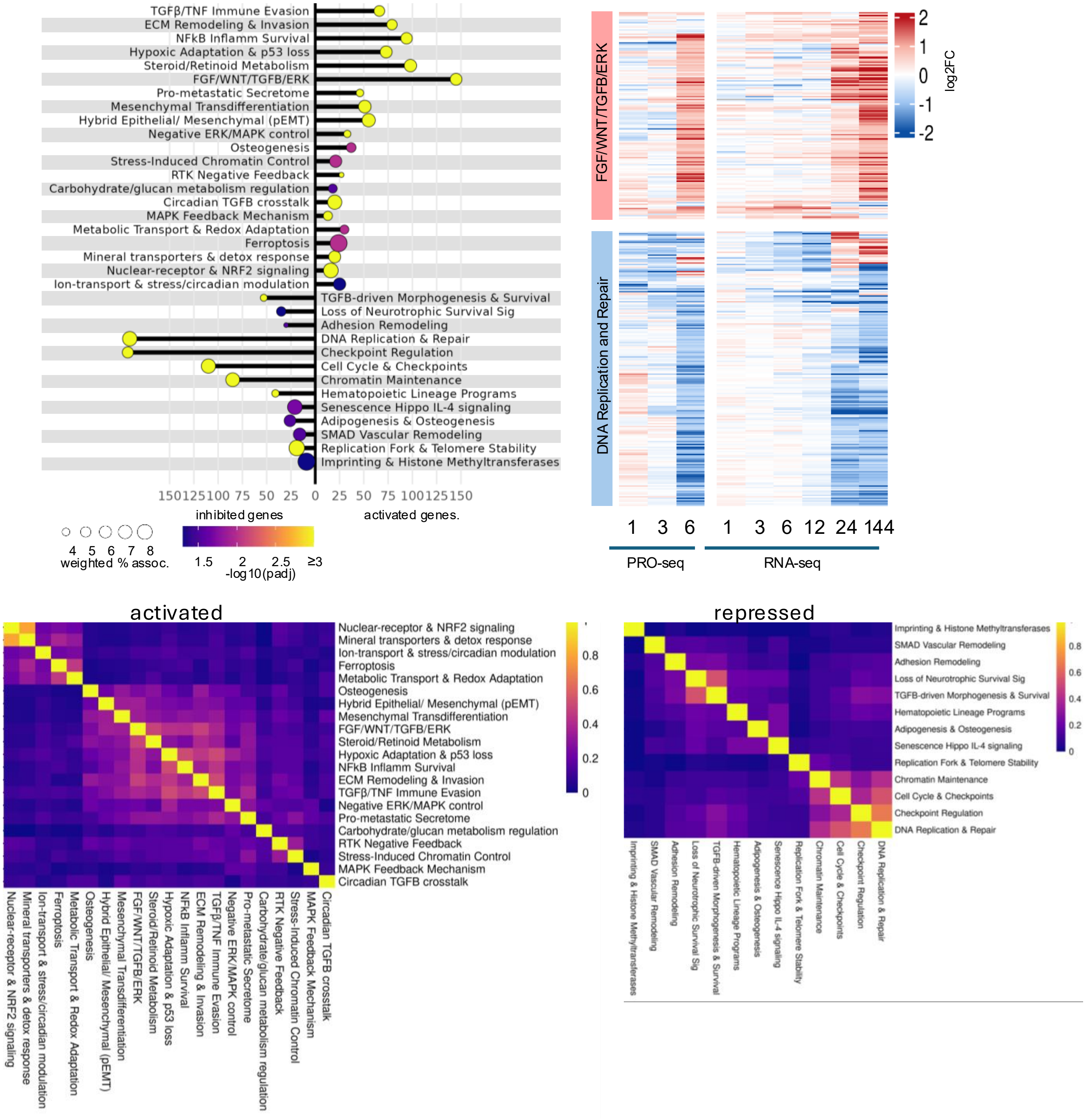


**Supplementary Figure 4. Functional organized network of biological terms of DIG/ DEGs upon RUNX1 loss.** A) Functionally grouped network ClueGo analysis for DIG/ DEG upon RUNX1 ablation. Color scale indicates padj values and bubble size indicates weighted percent association defined pathways. Heatmap of row normalized z-score values of select genes associated with B) FGF signaling and C) DNA damage response. Jaccard similarity matrix of D) upregulated and E) downregulated genes.

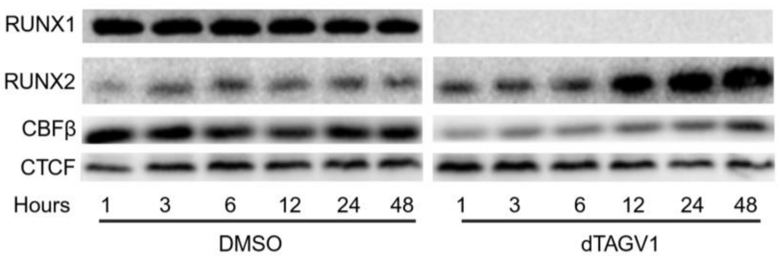


**Supplementary Figure 5. RUNX2 protein expression increases after 12hrs post RUNX1 ablation.** Western blot demonstrating that RUNX2 increases in protein expression but only after 12hrs of dTAGV1 treatment well after changes in gene expression. Exposure time for RUNX2 was significantly longer than RUNX1 indicating that RUNX2 expression is qualitatively less than RUNX1.
